# A Spatiotemporal Molecular Atlas of the Ovulating Mouse Ovary

**DOI:** 10.1101/2023.08.21.554210

**Authors:** Madhav Mantri, Hanxue Hannah Zhang, Emmanuel Spanos, Yi A Ren, Iwijn De Vlaminck

**Affiliations:** Nancy E. and Peter C. Meinig School of Biomedical Engineering, Cornell University, Ithaca, New York; Department of Animal Science, Cornell University, Ithaca, New York

**Author notes:** These authors contributed equally (Yi A Ren, Iwijn De Vlaminck).

## Abstract

Ovulation is essential for reproductive success, yet the underlying cellular and molecular mechanisms are far from clear. Here, we applied high-resolution spatiotemporal transcriptomics to map out cell-type- and ovulation-stage-specific molecular programs as function of time during follicle maturation and ovulation in mice. Our analysis revealed dynamic molecular transitions within granulosa cell types that occur in tight coordination with mesenchymal cell proliferation. We identified new molecular markers for the emerging cumulus cell fate during the preantral-to-antral transition. We describe transcriptional programs that respond rapidly to ovulation stimulation and those associated with follicle rupture, highlighting the prominent roles of apoptotic and metabolic pathways during the final stages of follicle maturation. We further report stage-specific oocyte-cumulus cell interactions and diverging molecular differentiation in follicles approaching ovulation. Collectively, this study provides insights into the cellular and molecular processes that regulate mouse ovarian follicle maturation and ovulation with important implications for advancing therapeutic strategies in reproductive medicine.

## INTRODUCTION

Ovulation is a complex process that involves extensive molecular, cellular and structural changes in distinct compartments in ovarian follicles, which culminate in follicle wall rupture and release of oocytes for fertilization. Successful ovulation is predicated on precise coordination and interaction between cell types within the ovary, both temporally and spatially. Temporally, the gene expression processes that are important for ovulation are regulated dynamically on a time scale of minutes to hours^1^. Spatially, the formation of follicle rupture sites requires spatially restricted tissue remodeling between granulosa and mesenchymal cells in the surrounding stromal tissues^2^. The cellular and molecular mechanisms responsible for controlling this temporal and spatial specificity of follicle rupture, however, remain poorly understood.

Single-cell RNA-sequencing has been used extensively to characterize the diversity of cell types and phenotypes in the ovary^3–6^. Yet, single-cell RNA sequencing requires tissue dissociation and therefore leads to the loss of spatial information regarding cellular niches, cell-cell interactions and the interplay between morphological changes and cellular changes in tissues^7,8^. Spatial RNA sequencing has been applied recently to study the cellular organization within the ovary, but at limited spatial resolution and without incorporating temporal analyses^9,10^. Here, we address these challenges by implementing spatial RNA sequencing at 10 μm spatial resolution and by profiling gene expression at eight well defined stages preceding ovulation in mice. To induce ovulation, we used exogenous gonadotropin stimulation in immature mice, a widely used model to study the molecular and cellular mechanisms that regulate ovulation. To support our analysis, we developed new analytical approaches to study cell-cell interactions in specific spatial niches within ovarian follicles. Altogether, our work leads to a highly detailed spatiotemporal atlas of ovulation in mice and provides insights into how ovarian follicle maturation and rupture are regulated in time and space.

## RESULTS

### High-resolution spatial transcriptomics of murine ovaries undergoing ovulation

To investigate the spatiotemporal regulation of ovarian follicle maturation and ovulation in mice, we implemented high-resolution spatial gene expression profiling on ten ovaries, collected at eight well-defined time points between ovulation induction and follicle rupture (Seeker, Curio Biosciences, **Methods, Fig. 1A & 1B**). Following an initial quality control — removing bead transcriptomes with fewer than 100 unique molecules detected and those not located directly under the tissue —we analyzed 121,536 spatial transcriptomes (**Supp. Fig. 1A-1C**). We assigned the most likely cell type to each spatial transcriptome via deconvolution using a scRNA-seq reference (**Methods**). The spatial transcriptomes represented seven broad cell types: granulosa cells, mesenchymal cells, oocytes, endothelial cells, epithelial cells, immune cells, and erythrocytes (**Fig. 1C, Supp. Fig. 1D**). As each bead used in spatial transcriptomics measures 10 μm, this dataset provides the first near single-cell resolution, whole-genome spatiotemporal transcriptomic atlas of immature and ovulating ovaries.

**Figure 1.**
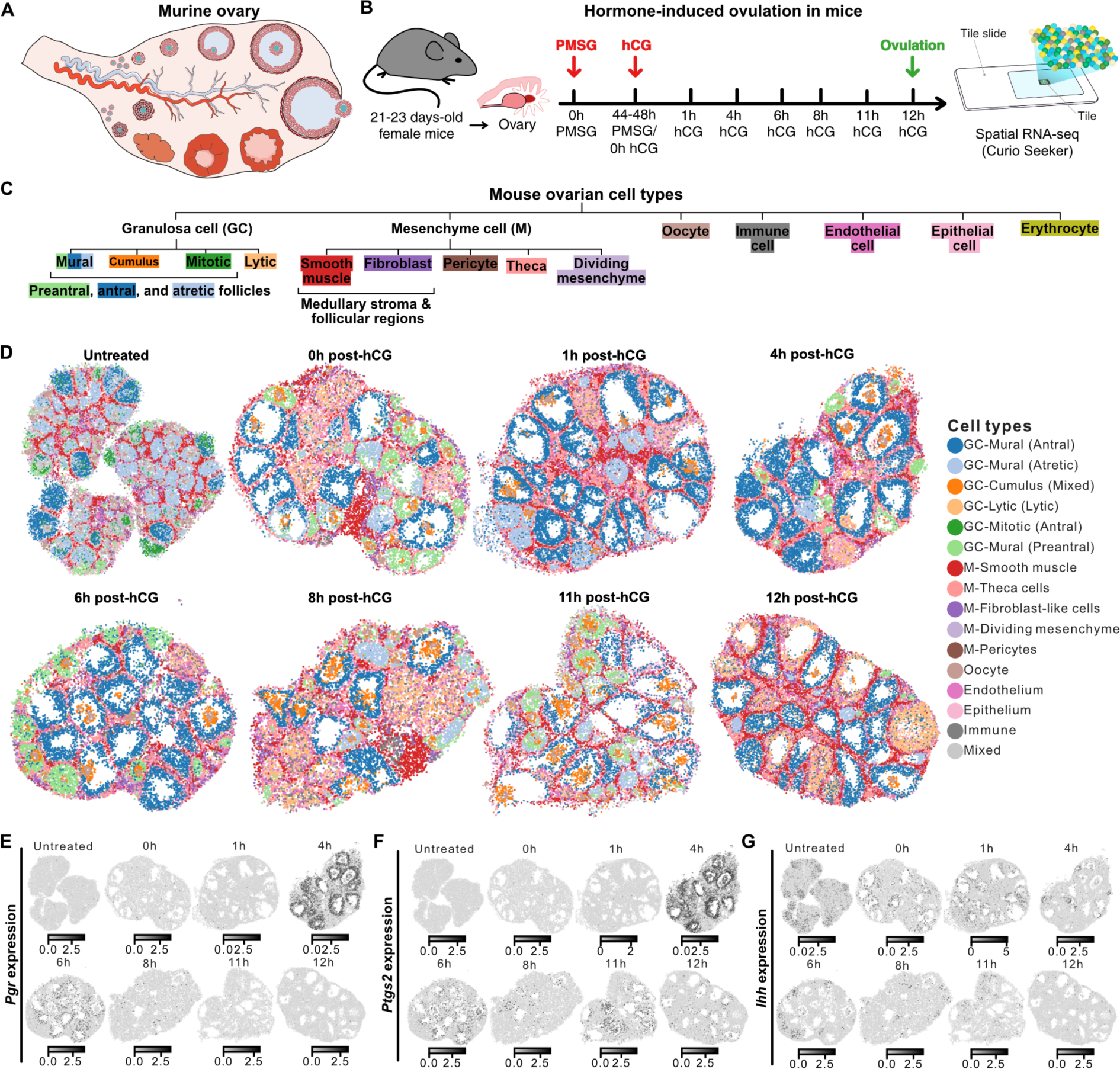
Spatial transcriptomics profiling of murine ovaries undergoing hormone-induced ovulation. **A)** Illustration depicting the internal components of the mouse ovary. **B)** Workflow schematic outlining the steps followed in the experiment. PMSG = pregnant mare serum gonadotropin; hCG = human chorionic gonadotropin. Spatial transcriptomics was conducted on tissue sections from three untreated immature ovaries and seven ovaries undergoing hormone-controlled ovulation. **C)** Tree diagram showing multi-level hierarchy of different cell types within the ovary. The first level of the tree represents the broad cell type labels predicted for Slide-seq beads using scRNA-seq data from cycling mouse ovaries. The leaves of the tree represent the fine cell type labels assigned using canonical markers after unsupervised clustering of bead transcriptomes in individual samples. **D)** Spatial transcriptomics maps of three immature ovaries and seven representative ovaries undergoing hormone-controlled ovulation colored by fine-grained cell type labels. **E-G)** Spatial transcriptomics maps showing expression of *Pgr*, *Ptgs2*, and *Ihh* over eight time points before and after ovulation stimulation.

To refine the classification of granulosa cell types, we used additional clustering and analysis of expression of canonical cell-type specific markers in conjunction with observed spatial location within the tissue (**Supp. Table 1, Supp. Fig. 1C & 2A, Methods**). This allowed us to identify mural cells located along the follicular walls, and cumulus cells surrounding the oocytes at each stage during ovulation (**Fig. 1D, Supp. Fig. 2A & 2B**). We further identified four types of mural granulosa cells with distinct transcriptomic profiles that are linked to follicle state (preantral, antral, atretic, and lytic follicles, **Fig. 1D, Supp. Fig. 2A & 2B**). Additionally, we refined cell type labels for beads within the mesenchyme via deconvolution using a scRNA-seq reference^3^, and identified smooth muscle cells, fibroblast-like cells, theca cells, dividing mesenchyme cells, and pericytes (**Fig. 1C, 1D, Supp. Fig. 2C & 2D**). Mesenchymal smooth muscle and fibroblast-like cells were visible in some of the ovarian tissue sections both in the medullary stroma (**Supp Fig. 2C & 2D**) and associated with growing follicles (**Supp Fig. 2C**). To enable the characterization of cell-cell interactions and molecular changes specific to defined stages of follicle maturation, we manually segmented the 335 follicles represented in the ten ovary tissue sections (**Supp Fig. 3A-C**). Spatiotemporal maps for the transcripts of *Pgr*, *Ptgs2* and *Ihh* demonstrated patterns of expression consistent with previous reports^11–14^ (**Fig. 1E, 1F and 1G**). Collectively, we generated a spatially resolved hierarchical map of the cell types and their phenotypes in immature and preovulatory mouse ovaries during hormone-induced ovulation.

**Figure 2.**
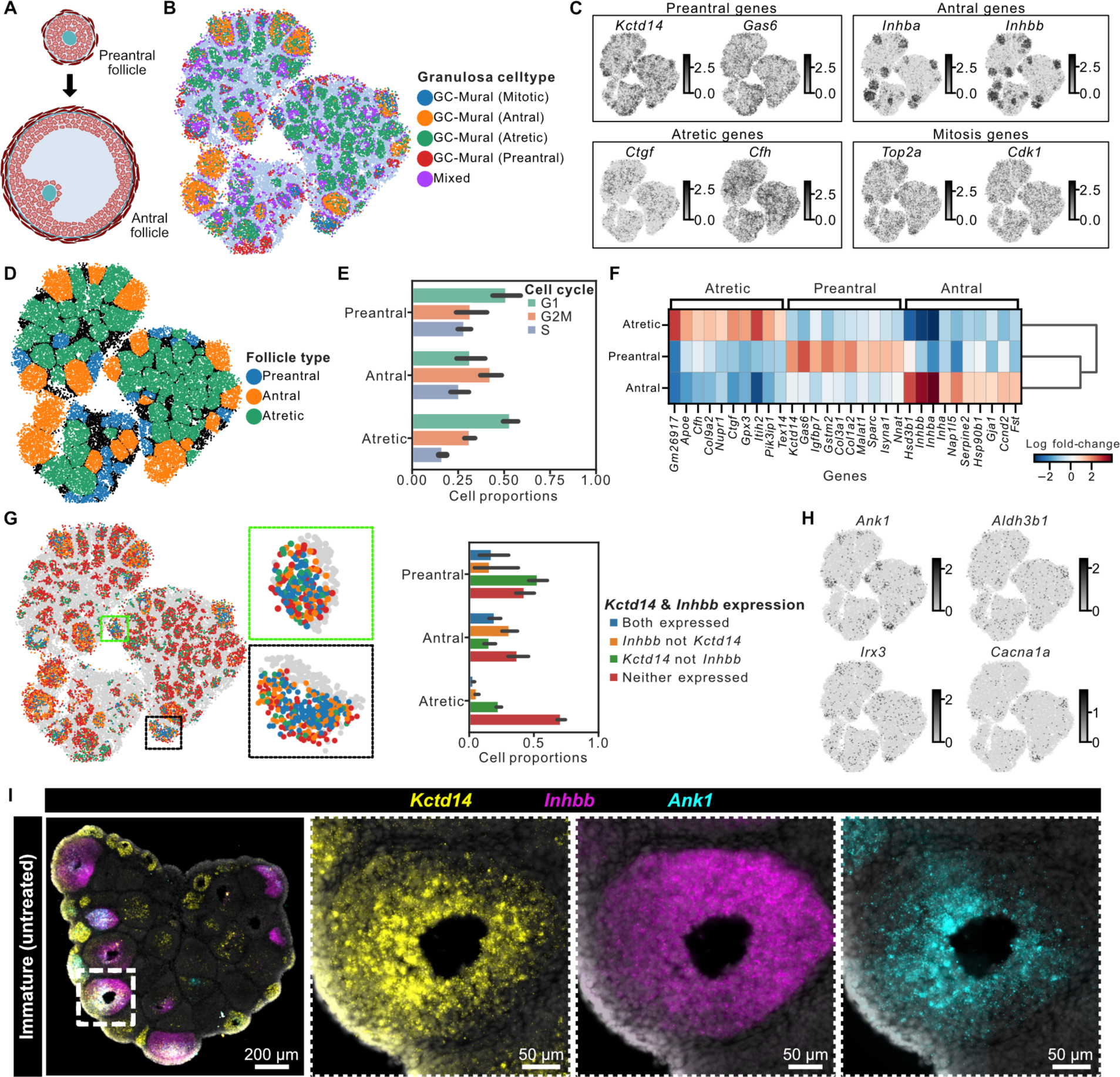
Spatial and molecular differentiation of distinct granulosa cell layers emerging during prenatal-to-antral transition. **A)** Illustration showing the structural transition from preantral to antral ovarian follicles. **B)** Spatial transcriptomics maps of three untreated immature ovaries colored by granulosa cell type. **C)** Spatial expression of three immature untreated ovaries showing the expression of previously reported and novel gene markers for preantral, antral, atretic, and mitotic follicles. **D)** Spatial transcriptomics maps of immature ovaries colored by follicle type as determined by gene expression of mural granulosa cells within the follicles. **E)** Bar plot showing proportion of cells in various cell cycle phases across different follicle types. **F)** Heatmap showing differentially expressed genes in mural granulosa cells across different follicle types. **G)** Spatial transcriptomics maps showing spatial organization for the expression of preantral follicle marker, *Kctd14*; and antral follicle marker, *Inhbb*. **H)** Spatial transcriptomics maps showing the expression of genes specific to granulosa cells in follicles transitioning from preantral to antral stage: *Ank1*, *Aldh3b1*, *Irx3*, and *Cacna1a*. **I)** Multiplexed RNA FISH staining for preantral marker *Kctd14* (yellow), antral marker *Inhbb* (magenta), and *Ank1* gene (cyan) on untreated immature ovary. The dotted boxes show zoomed-in images of a representative antral follicle. Representative images are from five biological replicates.

**Figure 3.**
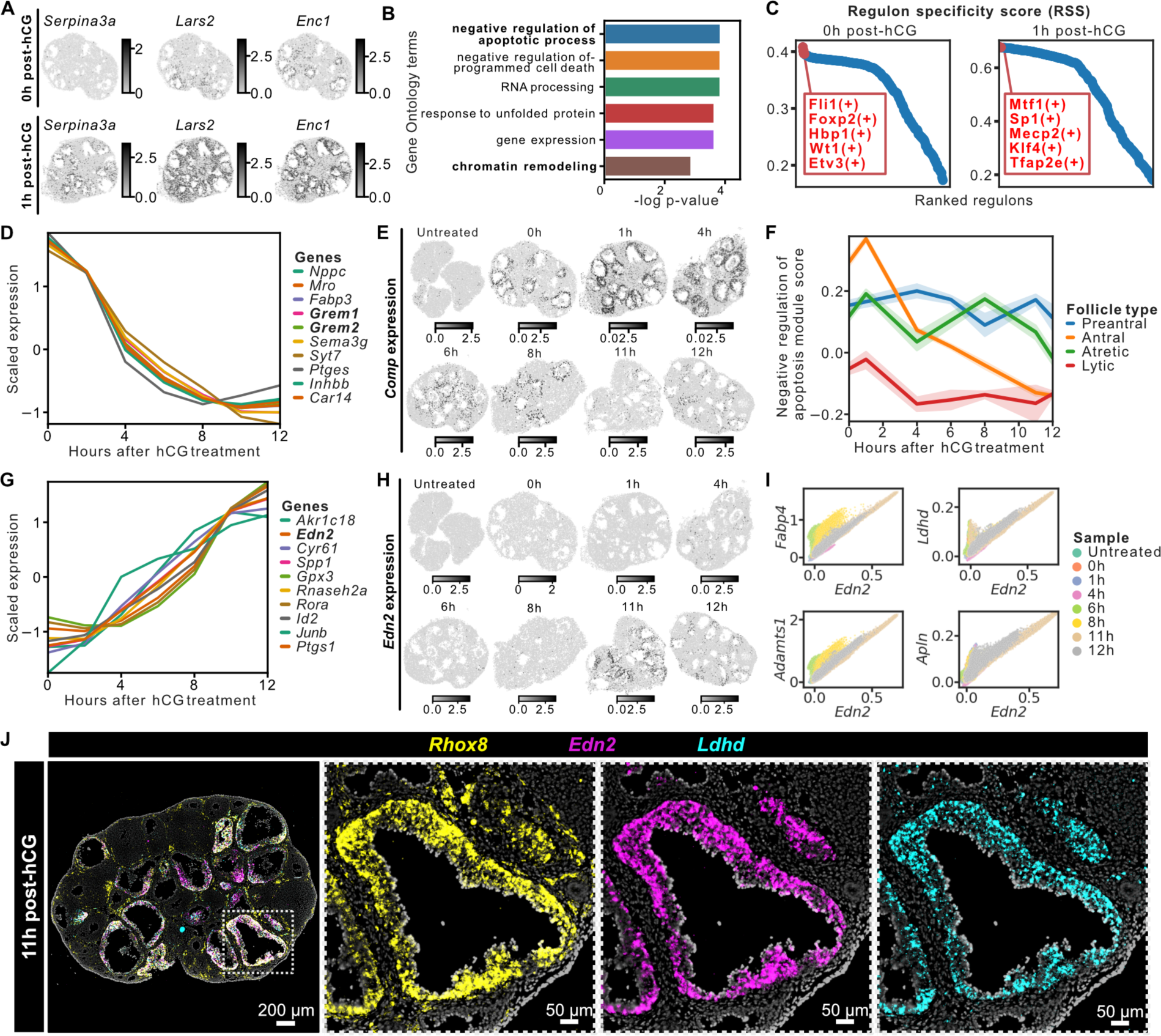
Spatiotemporal response of mural granulosa cells to ovulation induction. **A)** Spatial maps showing genes upregulated in mural granulosa cells one hour after PMSG and hCG hormone treatment. **B)** Bar plot showing the top six gene ontology (GO) terms for genes upregulated one hour after hCG treatment. **C)** Rank for gene regulatory networks, referred to as regulons, in mural granulosa cells based on regulon specificity score (RSS) one-hour post-hCG treatment. Top five most specific regulons at 0h and 1h post-hCG is labeled in red. **D)** Line plot showing temporal trends of genes that monotonously decreased with time post-hCG in mural granulosa cells of preovulatory follicles. **E)** Spatial transcriptomics map showing the temporal expression of *Comp* in immature and preovulatory ovaries. **F)** Spatial transcriptomics map showing the temporal gene module score for negative regulation of apoptosis in untreated immature and preovulatory ovaries. **G)** Line plot showing temporal trends of genes that monotonously increased with time post-hCG in mural granulosa cells from preovulatory follicles. **H)** Spatial transcriptomics map showing the temporal expression of *Edn2* in untreated immature and preovulatory ovaries. **I)** Scatter plots showing expression of four genes that correlate with the expression of *Edn2* in mural granulosa cells. Each point in the scatter plot represents a spatial transcriptome. **J)** Multiplexed RNA FISH staining for granulosa cell marker *Rhox8* (yellow), rupture-associated marker *Edn2* (magenta), and metabolic activity gene *Ldhd* (cyan) in preovulatory follicles at 11h post-hCG. The dotted boxes show zoomed-in images of a representative preovulatory follicle. Representative images are from three biological replicates.

### Spatial delineation and novel molecular markers of granulosa cells associated with preantral-to-antral transition

We first analyzed granulosa cells within the immature ovaries to map their molecular changes during folliculogenesis prior to ovulation induction (**Fig. 2A**). Unsupervised clustering identified four distinct granulosa cell types: ***i)*** granulosa with preantral phenotype expressing markers *Kctd14* and *Gas6*, ***ii)*** granulosa with early antral phenotype expressing markers *Inhba* and *Inhbb*, ***iii)*** granulosa with atretic phenotype expressing markers *Ctgf* and *Cfh*, ***iv)*** and granulosa cells with mitotic phenotype expressing *Top2a* and *Cdk1* (**Fig. 2B & 2C**). We used the phenotype of granulosa cells to classify follicles as preantral (*Kctd14*+ granulosa cells), antral (*Inhbb*+ granulosa cells) or atretic (*Cfh*+ granulosa cells, **Fig. 2D**). The majority of follicles in immature ovaries were atretic, and most antral follicles were located along the edge of the ovary. Consistent with the absence of antrum observed in H&E staining, preantral follicles mapped by spatial transcriptomics were completely filled with granulosa cells (**Supp. Fig. 1C**). Notably, mitotic granulosa cells in G2M phase were more prevalent in antral follicles, consistent with the observation of *Top2a* and *Cdk1* expression in these follicles, and in line with the expected growth of antral follicles (**Fig. 2E**). Interestingly, in atretic follicles, while the outer layers of granulosa cells expressed markers of degeneration, the inner layers of granulosa cells expressed mitotic markers (**Supp. Fig. 4A & 4B**). Last, we used the follicle segmentation and classification to identify novel markers of granulosa cells associated with follicle maturation stage. *Gas6*, *Gstm2*, *Isyna1* were enriched in preantral follicles; *Nap1l5* in antral follicles; and *Gm26917*, *Itih2* in atretic follicles (**Fig. 2F**).

The transition between preantral and antral follicle is a crucial stage of follicle maturation. We overlaid markers of preantral (*Kctd14*) and antral (*Inhbb*) follicle phenotypes, which revealed intricate spatial patterns of granulosa cell differentiation markers during the preantral-to-antral follicle transition (**Fig. 2G**). Preantral and early antral follicles showed an enrichment of a *Inhbb*+*Kctd14*+ cell population in the innermost layers of granulosa cells (boxed follicles: upper, preantral; lower, early antral. Histology staining of these follicles is shown in **Supp. Fig. 1C**. In the same follicles, the *Inhbb*+*Kctd14*-granulosa cells surrounded the double-positive cells. As antral follicles grew bigger, the innermost granulosa cells retained the expression of both *Inhbb* and *Kctd14* while the population of *Inhbb*+*Kctd14*-granulosa cells expanded. In preovulatory follicles, we found that mural granulosa cells expressed only *Inhbb*, while cumulus cells expressed both *Inhbb* and *Kctd14* (**Supp. Fig. 4C-4E**). Together, these observations strongly suggest that *Inhbb*+*Ktcd14*-cells along the follicle wall are precursors of mural granulosa cells, and that the *Inhbb*+*Kctd14*+ cells in the center of small antral follicles are precursors of cumulus cells, which corroborates with a previous report on the expression of *Kctd14* in preantral follicles and cumulus cells^3^. In summary, these observations revealed distinct layers of granulosa cells emerging and expanding during preantral-to-antral transition.

To delve deeper into the molecular signatures underlying the transition between preantral to antral follicles, we performed differential gene expression analysis and found *Ank1*, *Aldh3b1*, *Irx3*, and *Cacna1a* upregulated in *Inhbb*+*Kctd14*+ double-positive granulosa cells (**Fig. 2H, Supp. Fig. 4F**). We performed multiplexed Fluorescence In Situ Hybridization (FISH) to visualize *Ank1* expression in ovaries from untreated mice, and found that *Ank1* (ankyrin 1 that regulates cell membrane stability and shape) was indeed expressed in the *Inhbb*+*Kctd14*+ double-positive granulosa cells surrounding oocytes (**Fig. 2I**). Intriguingly, in preantral follicles where all granulosa cells expressed *Kctd14* but not *Inhbb*, *Ank1* was enriched in granulosa cells surrounding the oocytes towards the center of the follicles; while in preovulatory follicles, *Ank1* expression is retained in cumulus cells but not mural granulosa cells (**Supp. Fig. 4G & 4H**). We suggest that *Ank1*, *Aldh3b1*, *Irx3* and *Cacna1a* are early molecular markers for further differentiation of the cumulus cell fate, and that their induction of expression is associated with the preantral-to-antral follicle transition. Taken together, our findings delineate molecular and spatial changes in distinct granulosa cell layers and identified novel molecular regulators during the crucial preantral-to-antral transition.

### Spatiotemporal response of mural granulosa cells to ovulation stimulation

To delineate molecular changes driving follicle maturation and ultimately ovulation, we first analyzed the rapid transcriptomic response of mural granulosa cells to hCG treatment that mimics the preovulatory luteinizing hormone surge. Differential gene expression analysis identified 77 genes with significantly upregulated transcript levels (log fold change > 1.0, adjusted p-value < 0.01) as early as one-hour post-hCG treatment (0h post-hCG vs. 1h post-hCG, **Supp. Fig. 5A & Fig. 3A**), including *Serpin3a*, *Lars2*, and *Enc1.* As expected, they were predominantly expressed in mural granulosa cells of antral follicles (**Fig. 3A**). Gene ontology analysis of genes upregulated in mural granulosa cells one-hour post-hCG revealed an enrichment of ontology terms representing negative regulation of apoptosis and chromatin remodeling (**Fig. 3B**). Consistent with previous reports that hCG/LH inhibits granulosa cell apoptosis^15–17^, our analysis revealed an acute induction of transcripts that negatively regulate apoptotic processes and of cell cycle regulator *Ccnd2* in preovulatory follicles at 0-1h post-hCG^18,19^ (**Supp. Fig. 5B & 5C**). Regulatory network analysis on spatial transcriptomes of mural granulosa cells further revealed regulons that are responsive to hCG treatment (**Methods**), including *Mtf1*(+), *Tfap2e*(+), *Sp1*(+), *Klf4*(+), and *Mecp2*(+) (**Fig. 3C**). *Tfap2e*, *Sp1* and *Klf4* regulons have been implicated in transcriptional regulation of steroidogenesis^20^. Consistent with previous work^21^, we also observed in mural granulosa cells of preovulatory follicles a sharp increase in transcript levels of *Cyp19a1* between 0-1h post-hCG, before they decreased to base levels after 4h post-hCG (**Supp. Fig. 5D**). Interestingly, expression of *Cyp17a1* in theca cells was also initially upregulated and then decreased to base levels with similar temporal dynamics as *Cyp19a1* (**Supp. Fig. 5E**). These observations support a temporally coordinated increase in steroidogenic activities between mural granulosa cells and theca cells as a rapid response to hCG stimulation.

We further clustered genes into groups based on their temporal expression patterns in mural granulosa cells (**Methods, Supp. Fig. 5F**). Genes that exhibited a gross trend of gradual decrease in expression following hormone treatment included *Nppc*, an important regulator of oocyte meiotic arrest^22^, and those involved in apoptosis regulation, *Grem1*, *Grem2*^23^, and *Comp* (**Fig. 3D & 3E, Supp. Fig. 5G**). *Comp* (also known as *thrombospondin-5*, a non-collagenous extracellular matrix protein that suppresses apoptosis), was expressed in mural granulosa cells and cumulus cells of small antral follicles in the immature ovaries, and in large antral follicles of preovulatory ovaries, with its expression increasing abruptly at 0-1h post-hCG treatment but gradually declining after 1h post-hCG treatment (**Fig. 3E**). We calculated and visualized a module score for a panel of 12 genes associated with negative regulation of apoptosis and found this score first increased between 0-1h post-hCG, then sharply decreased in mural granulosa cells of the preovulatory follicles in contrast to preantral and atretic follicles (**Fig. 3F, Supp. Fig. 5H**). The initial increase of this gene module score is consistent with an upregulation of genes that negatively regulate apoptosis (**Fig. 3B**). We also found that, between 4h and 8h post-hCG, while this module score gradually decreased in mural granulosa cells of preovulatory follicles, it remained at appreciable levels in cumulus cells until immediately before ovulation (**Supp. Fig. 5I**). These observations support an important role of apoptosis regulation in the final stage of follicle maturation and preparation for ovulation.

Among genes that exhibit a continuous increase in expression in response to hormone treatment, *Edn2* emerged as one of the most variable genes. *Edn2* was expressed in mural granulosa cells of the preovulatory follicles immediately prior to ovulation (**Fig. 3G and 3H**). *Edn2* is known for its crucial role in ovulation^24,25^. To uncover potential novel regulators of follicle rupture and the transition to luteal phase, we searched for genes that are spatially and temporally associated with *Edn2* in preovulatory follicles (**Methods**). *Adamts1* was among these genes, consistent with previous reports of its important role in ovulation^12,26,27^ (**Fig. 3I**). We also identified a gradual increase in the expression of *Apln* in mouse mural granulosa cells, consistent with the role of *Apln* in ovarian steroidogenesis as reported in several species other than mice^28,29^ (**Fig. 3I**). *Cyr61* (also known as *Ccn1*) promotes adhesion of endothelial cells and upregulates angiogenic genes^30^, both important processes for the formation of corpus luteum. *Gpx3* codes for glutathione peroxidase, where glutathione protects cells from oxidative stress and is associated with fertility^31^. Notably, among genes that shared a similar expression pattern as *Edn2*, we found an enrichment of genes regulating cell metabolism, including *Akr1c18*^32^ (more commonly known as *20alpha-HSD*; metabolizes progesterone to its inactive form), *Rora* (an orphan nuclear receptor regulating cell differentiation, circadian gene expression and cell metabolism), *Ldhd* (lactate dehydrogenase D; catabolizes D-lactate to pyruvate), and *Fabp4*^33^ (regulates long-chain fatty transport and metabolism), which is consistent with the results from gene ontology enrichment analysis (**Supp. Fig. 5K & 5L**). Multiplexed FISH demonstrated that *Ldhd* was not expressed in the ovary at 0h post-hCG but co-expressed with *Rhox8* and *Edn2* in mural granulosa cells shortly before ovulation (**Fig 3J and Supp. Fig. 5J**). Together, these observations reveal temporally coordinated changes of metabolic activities and follicle rupture in mural granulosa cells during hormone-induced ovulation.

### Dynamic and temporally controlled oocyte-cumulus cell interactions in preovulatory follicles

The rupture of ovarian follicles at ovulation must be closely coordinated with meiotic maturation of oocytes to ensure successful fertilization^34^ (**Fig. 4A**). To gain insights into the molecular changes associated with oocyte maturation, we analyzed the transcriptomes of individual oocytes represented in the spatial transcriptomic data (**Fig. 4B**). We implemented stringent quality control (**Methods**) and produced pseudo-bulk transcriptomes specific to 83 whole oocytes (average number of beads per cell = 16, **Supp. Fig. 6A & 6B** and **Fig. 4B & 4C**). Visualization of the total number of detected genes within oocytes in response to hormone treatments revealed an initial increase in transcriptional diversity up to 6h post-hCG treatment, followed by a subsequent decrease (**Fig. 4D**). Clustering analysis of oocyte transcriptomes showed only subtle variations among oocytes from different follicle types and stages, with one notable exception: a well-defined cluster of oocytes obtained from ovaries at 11h and 12h post-hCG (**Supp. Fig. 6B & 6C**). We observed an upregulation of *Btg4*, *C86187*, *Riok1*, and *Ccdc186* in these mature oocytes (**Fig. 4E**). Of functional interest to oocyte maturation and early embryogenesis, we also found in this cluster an enrichment of genes that regulate RNA activities (*Dnajc21*, *Zcchc8*, *Unc50* and *Rbm36*), and DNA demethylation (*Riok1* and *Tet3*). These results provide valuable insights into the coordination of ovulation and oocyte maturation.

**Figure 4.**
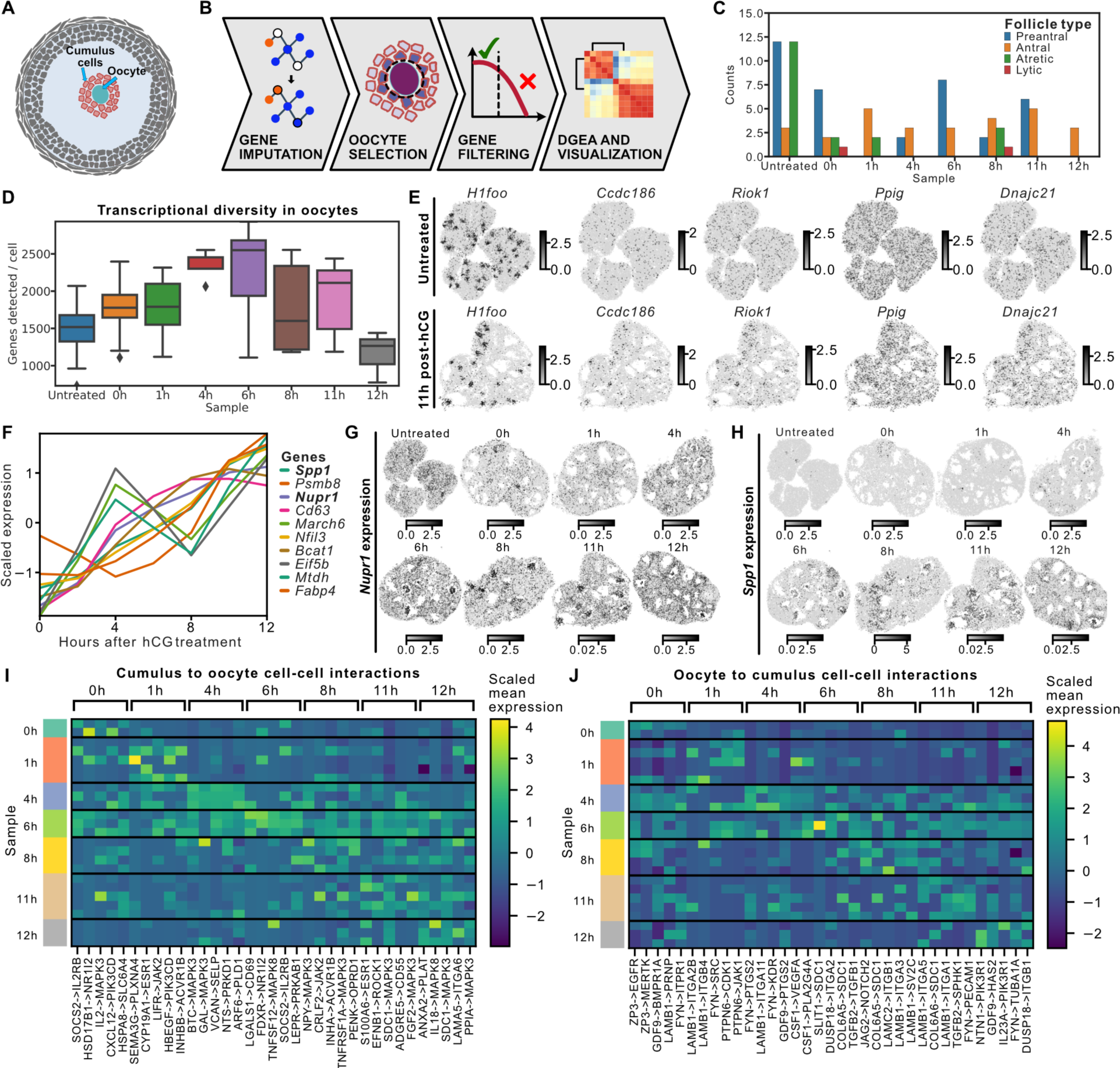
Dynamic and temporally controlled oocyte-cumulus cell interaction in preovulatory follicles. **A)** Illustration showing the structure of a preovulatory follicle with oocyte and surrounding cumulus cells. **B)** Analysis workflow for oocyte segmentation and characterization of oocyte heterogeneity across follicle types. **C)** Bar plot showing the counts of segmented oocytes in preovulatory ovaries at different stages across different follicle types. **D)** Box plot showing the number of genes detected per oocyte across time points. **E)** Spatial transcriptomics maps of immature and ovulating ovaries showing the expression of a pan-oocyte marker (*H1foo*) and four genes enriched in mature oocytes: *C86187*, *Riok1*, *Ppig*, and *Dnajc21*. **F)** Line plot showing temporal trends of genes that show a gradual increase in expression over time post-hCG in cumulus cells from preovulatory follicles. **G)** Spatial transcriptomics map showing the temporal expression of *Nupr1* in immature and preovulatory ovaries. **H)** Spatial transcriptomics map showing the temporal expression of *Spp1* in immature and preovulatory ovaries. **I)** Heatmap showing the mean expression of top-five differentially expressed ligand-receptor genes expressed in cumulus cells and oocytes from preovulatory follicles, respectively. Each row represents one cumulus-oocyte-complex (COC). **K)** Heatmap showing the mean expression of top-five differentially expressed ligand-receptor genes expressed in oocytes and cumulus cells from preovulatory follicles, respectively. Each row represents one COC.

Cumulus cells provide the oocyte with crucial nutrients and undergo expansion in response to ovulation induction. We computed trends in the expression of highly variable genes in cumulus cells and clustered the genes into groups based on their temporal expression patterns (**Methods, Supp. Fig. 6D-6F**). In line with existing knowledge, we observed upregulation of *Ptx3*, which encodes pentraxin-3, a major component of the extracellular matrix of cumulus cells during expansion^35^ (**Supp. Fig. 6E & 6I**). Among genes that exhibited a sharp increase in expression between 0h and 4h post-hCG, *Star* and *Cyp11a1* emerged as the most variable gene (**Supp. Fig. 6G & 6H**), supporting previous findings on steroidogenic activities of cumulus cells after LH/hCG stimulation^31^. Among genes that display a more gradual increase in expression were *Nupr1*, which encodes a nuclear protein transcriptional regulator, and *Spp1*, which encodes secreted phosphoprotein that binds to the extracellular hydroxyapatite and is responsible for extracellular matrix organization (**Fig. 4F-G, Supp. Fig. 6J**).

The interaction between cumulus cells and oocytes is highly dynamic and context dependent. We performed bi-directional cell-cell signaling analysis for cumulus and oocyte cells within individual follicles at different time points between hCG stimulation and ovulation (**Methods**). We found time-specific cellular interactions from cumulus to oocyte post-hCG: INHBB->ACVR1B at 1h; BTC->MAPK3 and VCAN->SELP at 4h; NTS->PRKD1 at 4h and 6h; NPY->MAPK3 at 8h, FGF2->MAPK3 at 11h, and IL18->MAPK3 at 12h (**Fig. 4I**). We further identified novel ligand-receptor interactions, including SDC1->MAPK3 from 6h to 12h post-hCG and S100A6->ESR1 specifically enriched immediately before ovulation at 11h and 12h post-hCG (**Fig. 4I**). Similarly, we found novel communication pathways from oocytes to cumulus cells post-hCG: such as PTPN6->CDK1/JAK1 at 1h, FYN->PTGS2 at 4h, JAG2->NOTCH2 at 8h, and NTN1->PIK3R1 at 12h post-hCG (**Fig. 4J**). Collectively, these observations reveal new cellular interactions in the dynamic and temporally controlled cellular niche of cumulus cells and oocytes during the preovulatory stage.

### Spatiotemporally restricted cellular interactions and molecular programs during follicle maturation and rupture

Maturation and ovulation of ovarian follicles requires significant remodeling of the surrounding stromal tissues and acute formation of rupture sites on the ovary’s surface, facilitating the release of oocytes for fertilization. We quantified the proportions of different mesenchymal cell types and observed enrichment in the proportion of dividing mesenchymal cells, first in immature ovaries and then at 6h post-hCG (**Supp Fig. 7A**). Visual examination of dividing mesenchymal cells in the immature ovaries revealed their specific localization around antral follicles enriched in proliferating granulosa cells, which suggested coordinated proliferation of the mesenchyme and granulosa cells (**Fig. 5A and Supp Fig. 4B**). To understand the molecular basis of this coordination, we explored cell-cell communication between the dividing mesenchyme and mural granulosa cells of antral follicles in the immature ovaries (**Methods**). This analysis revealed INHA/INHBA->TGFBR3, INHA/INHBB->ACVR1, and LAMA1/2->RPSA among the most significantly enriched ligand-receptor pairs (**Fig. 5B**). The enrichment in the proportion of dividing mesenchyme at 6h post-hCG can be attributed to increased vascularization of stromal cell layers surrounding the preovulatory follicles as suggested by an increase in the proportion of endothelial cells as well as pericytes between 4h and 6h post-hCG (**Supp Fig. 7**). Overall, these observations pinpoint tight coordination between mesenchyme and mural granulosa cells during follicular maturation.

**Figure 5.**
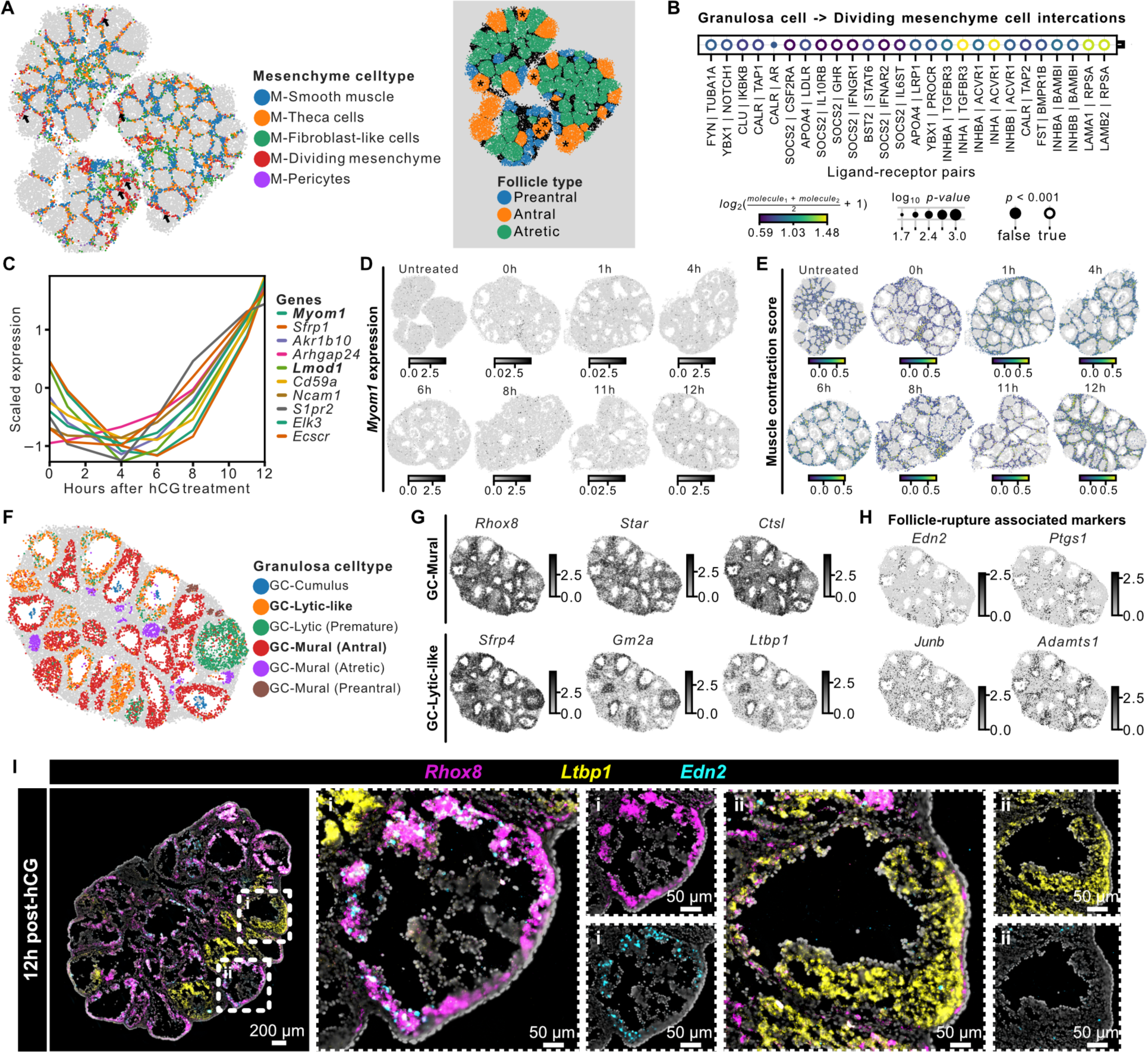
Spatially restricted molecular programs in ovarian stroma and periovulatory follicles. **A)** Spatial transcriptomics map of immature ovaries with mesenchymal transcriptomes colored by predicted mesenchymal cell types (left). Spatial transcriptomics map of immature ovaries colored by follicle type predicted based on the granulosa cell phenotypes (right). Arrows point to dividing mesenchyme cells in the untreated immature ovaries and asterisks on the follicle type map label the corresponding antral follicles. **B)** Dot plot showing ligand-receptor interaction between dividing mesenchymal cells and granulosa cells of antral follicles in immature ovaries. **C)** Line plot showing temporal trends of genes that display a sharp, late onset increase in expression post-hCG in mesenchymal cells. **D)** Spatial transcriptomics map showing the temporal expression of *Myom1* in immature and preovulatory ovaries. **E)** Spatial transcriptomics map showing the temporal gene module score for muscle contraction in immature and preovulatory ovaries. **F)** Spatial transcriptomics map of a preovulatory ovary at 12h post-hCG, with granulosa cell types colored by their transcriptomes. **G)** Spatial transcriptomics map of a preovulatory ovary at 12h post-hCG showing the expression of gene markers of mural granulosa cells (top) and lytic-like granulosa cells (bottom). **H)** Spatial transcriptomics map of a preovulatory ovary at 12h post-hCG showing the expression of follicle rupture-associated gene markers such as *Edn2*, *Ptgs1*, *Junb*, and *Adamts1*. **I)** Multiplexed RNA FISH staining for antral follicle associated granulosa cell marker *Rhox8* (yellow), lytic-like granulosa cell marker *Ltbp1* (cyan), and rupture-associated marker *Edn2* (magenta) in preovulatory follicles at 12h post-hCG. Representative images are from four biological replicates.

We identified three distinct time-dependent, mesenchyme-specific gene expression signatures: ***i)*** a sharp initial increase in the expression of genes associated with neurotrophic signaling from 0h to 4h post-hCG treatment (*Ntrk2*, *Bdnf*, and *Pdgfra); **ii)*** an intermediate increase in the expression of genes associated with extracellular structure organization between 4h and 8h post-hCG treatment (*Postn*, *Mmp2*, *Eln*, *Col5a2*, and *Tgfbi); **iii)*** and a sharp, late-onset increase in the expression of genes associated with myofibril assembly and muscle contraction at 8h post-hCG treatment (*Myom1*, *Lmod1*, *Pdgfrb*, **Fig. 5C and 5D, Supp Fig. 7**). We visualized a module score for genes associated with muscle contraction and found that this score is increased in the mesenchyme of preovulatory follicles after hCG treatment (**Fig. 5E**). This observation is in line with the reported role of muscle gene-expressing cells, in particular vascular smooth muscle cells, for ovulation^36–38^. Collectively, these results revealed distinct activities of the ovarian stroma during defined time windows preceding ovulation.

To gain new insight into follicle rupture and the transition to the luteal phase, we analyzed the granulosa cell transcriptomes at 11h and 12h post-hCG. While follicles appeared similar on histology immediately before follicle rupture at 12h post-hCG (**Supp. Fig. 1C**), we identified two distinct types of large ovulating follicles based on transcriptomic signatures of their granulosa cells (**Fig. 5F & 5G**). We asked whether these transcriptomic signatures are associated with the fate of the follicle to ovulate. And indeed, granulosa cells in one type of periovulatory follicles were enriched with transcripts of genes known to be crucial for ovulation, including *Edn2*, *Ptgs1*, *Junb*, and *Adamts1*^39–41^ (**Fig. 5H**). In contrast, transcript levels of these genes were low in the other type of follicles, which share a similar transcriptomic signature with granulosa cells in prematurely luteinized structures (lytic-like) (**Fig. 5G**). Multiplexed FISH of *Rhox8*, *Edn2*, and *Ltbp1* (latent transforming growth factor binding protein 1; a novel gene associated with of the lytic-like periovulatory follicles) confirmed that periovulatory follicles that express *Ltbp1* do not express *Edn2* (**Fig. 5I**) Hence, the molecular heterogeneity of periovulatory follicles may correspond to divergent follicle fates of ovulation versus premature luteinization. Interestingly, luteinization marker *Lhcgr* increased at 12h post-hCG in both types of periovulatory follicles, suggesting a transition stage right before ovulation (**Supp. Fig. 8A**). These findings highlight the rapid and intricate processes governing follicular fate determination immediately preceding ovulation.

## Discussion

We produced a spatiotemporal transcriptomic atlas using spatially resolved RNA-sequencing at near single-cell resolution to understand cell-type- and folliculogenesis-stage-specific mechanisms underlying mouse ovarian follicle maturation and ovulation. Our analysis revealed intricate spatial and molecular differentiation of distinct granulosa cell layers during the crucial preantral-to-antral transition^42,43^. We find that the transition toward the antral follicle state requires the coordinated expansion of *Inhbb*+*Kctd14*+ cells in the center layers surrounding the oocytes, and of *Inhbb*+*Kctd14*-mural granulosa cells surrounding the double-positive cells. We identified *Ank1* as a potential early marker for cumulus cell fate at the preantral stage. While *Kctd14* is ubiquitous in preantral follicles, we found that its expression is suppressed in the outer layers of granulosa cells of antral follicles. In contrast, we observed that *Ank1* is induced in inner layers of granulosa cells. This sharp spatial demarcation in molecular marker expression suggests that interactions between oocyte-derived factors and the spatial location of granulosa cells relative to the basement membrane dictates their differentiation and fate.

We further delineated cell-type-specific interaction mechanisms of ovarian cells. The changes that occur in the ovarian stroma and mesenchymal cells are important for follicle growth and rupture but challenging to study *in vivo*. Using spatial transcriptomics, we observed a surge in the proliferative activity of mesenchymal cells in early antral follicles which also had high proliferative activity of granulosa cells. These coordinated proliferative activities are possibly mediated by the ovarian local effect of inhibins and laminins. We observed a subsequent burst of mesenchymal cell proliferation at around 6h post-hCG in parallel with an enrichment of genes regulating extracellular matrix remodeling and an increase in the number of endothelial cells and pericytes. Transcriptomic changes in cumulus cells and oocytes in response to ovulation stimulation have been studied extensively^44–46^, but rarely without separating the two cell types *in vitro*, or at time points immediately preceding ovulation. We profiled oocyte-cumulus cell interaction mechanisms specific to defined post-hCG time points. Close to ovulation, we observed increased expression of *Nupr1*, *Spp1* and *S100a6* in cumulus cells, and an enrichment in the expression of genes that regulate RNA activity and DNA demethylation in oocytes.

In addition, we report gene regulatory programs at the follicle level both in rapid response to ovulation stimulation, and in association with the process of follicle rupture and transition to the luteal phase. Limited information was previously available on gene regulation during the first hour after hCG simulation and the last hour before ovulation. We found that during the first hour post-hCG, genes related to chromatin remodeling are upregulated, the expression of steroidogenic genes is increased in both mural granulosa cells and theca cells, and an anti-apoptotic program is induced, supporting the notion that coordinated cell survival and steroidogenic programs respond rapidly to hormone stimulation. In the hour leading up to follicle rupture, we observed genes such as *Akr1c18* and *Ldhd*, which modulate cell metabolism and steroidogenesis, show the same trend of increase as a gene module score of muscle contraction and genes associated with follicle rupture, including *Edn2* and *Adamts1*^12,24^. Immediately before follicle rupture, we find that periovulatory follicles exhibit acute luteinization and divergent heterogeneity, with a subpopulation of follicles poised to rupture. These observations highlight the dynamic biological processes that occur at the final stage of follicle maturation immediately before ovulation.

Given that regulators and pathways that are important for ovarian follicle maturation and ovulation are highly conserved between humans and mice, our findings, and data promise to translate into valuable insights for infertility management in humans.

## METHODS

### Ethical approval for animal experiments

All animal work was conducted ethically, conforming to the U.S. Public Health Service policy, and was approved by the Institutional Animal Care and Use Committee at Cornell University (IACUC approved protocol number 2019-0006). C57BL/6J mice were bred in-house for all experiments in the study. Mice were housed in 11.5” x 7.5” IVC Polycarb Shoebox Cage for the duration of the experiment. Temperature 68-77°F and humidity between 30 to 70% were maintained in the rodent room. Lights were turned ON from 5AM and OFF at 7PM in the rodent room.

### Hormone treatment for induction of ovulation

21-23 days old female mice with body weight of 10∼11 grams were treated with pregnant mare serum gonadotropin (PMSG) and human chorionic gonadotropin (hCG) to induce ovulation. Untreated mice of the same age were used as immature controls. Mice were first administered 5 IU PMSG (BioVendor, RP1782725000) followed by 5 IU hCG 48h later (EMD Millipore Corporation, #230734-1MG) in 100µl 0.1% BSA-PBS per mouse via intraperitoneal injection using a 27G insulin syringe. Three immature and seven preovulatory mice across eight different time points, namely untreated immature, 0h, 1h, 4h, 6h, 8h, 11h, and 12h post-hCG treatment, were used for spatial transcriptomics experiments. To evaluate the response of mice to hormone treatment, RNA extraction followed by reverse transcription (RT) and quantitative polymerase chain reaction (qPCR) was performed on one ovary from every mouse for a panel of selected genes. The other ovaries from the mice selected based on RT-qPCR results were used for spatial transcriptomics experiments.

### Sample preparation for Curio spatial transcriptomics

Whole ovaries were isolated using aseptic technique and placed in ice cold and sterile 1X Phosphate Buffered Saline (1X PBS, without calcium and magnesium chloride; Gibco 10010023). Blood and other contamination were carefully removed by washing the tissues with fresh 1X PBS. Fresh tissues were immediately embedded in Optimal Cutting Compound (OCT) media (SAKURA 25608-930) and frozen directly on dry ice for spatial transcriptomics experiments. The tissue blocks were cut into 10 µm sections using Thermo Scientific CryoStar NX50 cryostat and mounted on a Seeker Tiles (Curio Bioscience) for spatial transcriptomics experiments.

### Slide-seq spatial transcriptomics library preparation

Slide-seq spatial transcriptomics^47,48^ experiment was performed using the Curio Seeker Kit (Curio Bioscience) according to manufacturer’s instructions. Briefly, tissue sections from fresh-frozen murine ovaries were mounted on a 3mm x 3mm spatially indexed bead surface (Curio Seeker Kit, Curio Bioscience). After RNA hybridization and reverse transcription, the tissue sections were digested, and the beads were removed from the glass tile and resuspended. Second strand synthesis was then performed by semi-random priming followed by cDNA amplification.Sequencing libraries were then prepared using the Nextera XT DNA sample preparation kit. The libraries were pooled and sequenced on an Illumina NextSeq 2K (P3 flow cell) using the 100-cycle kit (Read 1 = 50 bp, Read 2 = 72 bp, Index 1 = 8 bp, and Index 2 = 8 bp). The sequencing data was aligned to the mouse genome (assembly: GRCm38) using the STAR Solo^49^ (version=2.7.9a) pipeline to derive a feature x bead barcode expression matrix. Bead barcode location files for the corresponding tiles were provided by Curio Bioscience.

### Slide-seq data preprocessing for quality control and smear removal

Slide-seq count matrix and the position information for every bead barcode were loaded into an AnnData object using scanpy^50^ (v1.9.1). After filtering out the beads with less than 100 transcripts detected, we log-normalized the Slide-seq expression data and computed principal components using highly variable genes. The transcriptomes were then clustered using the Leiden clustering algorithm and visualized on spatial maps. Cluster maps were then compared to Hematoxylin and Eosin (H&E) stained images of consecutive tissue sections, and clusters representing beads not located under the tissue were manually removed from individual Slide-seq datasets. Spatial distances between all pairs of beads within individual samples were then used to filter out beads with less than 10 beads within 100 µm distance. Individual samples were then reprocessed, and an additional round of filtering was performed to remove clusters of beads not located under the tissue. After filtering, individual datasets were processed and used for cell type labeling.

### Multi-level cell type label assignment for spatial transcriptomics datasets

scRNA-seq dataset from cycling murine ovaries^3^ was used as a reference to perform a multi-level deconvolution of the spatial transcriptomics datasets. First, cell2location^51^ (v0.1.3) was used to deconvolve spatial transcriptomes and broad cell type labels were assigned to each bead. Genes in the reference were filtered with cell_count_cutoff=5, cell_percent_cutoff=0.03, and nonz_mean_cutoff=1.12 to select for highly expressed markers of rare cell types while removing most uninformative genes. Cell type signatures were determined using NB regression and used for spatial mapping of scRNA-seq broad cell type labels on Slide-seq data with hyperparameters N_cells_per_location=1 and detection_alpha=20. Additional round of clustering was performed on beads labeled as granulosa cells within individual datasets and differential gene expression analysis in conjunction with observed spatial location within the tissue was used to label cumulus cells and four distinct types of mural granulosa cells with distinct transcriptomic profiles that are linked to follicle state (preantral, antral, atretic, and lytic follicles). Additionally, another round of deconvolution using a scRNA-seq reference was performed to refine cell type labels for beads classified as mesenchyme. Cell type signatures were determined using NB regression and used for spatial mapping of scRNA-seq fine cell type labels on Slide-seq data with hyperparameters N_cells_per_location=1 and detection_alpha=20. Fine cell type labels were assigned to mesenchymal cells located in both cortical and medullary stroma of the ovary tissue sections. All celltype labels were integrated together for visualization and analysis in the scanpy package.

### Follicle segmentation and follicle state assignment in Slide-seq datasets

Individual Slide-seq datasets were read using squidpy^52^ package (v1.2.3) and visualized in the interactive Napari image viewer^53^ (v0.4.15). Broad cell type labels assigned to individual beads were visualized on spatial maps and polygon shapes were drawn in the mesenchyme separating the follicles to manually segment 335 follicles across eight Slide-seq datasets. The beads within individual segmented follicles were assigned a unique follicle-ID for follicle level analysis and visualization. Each follicle was assigned a follicle state (preantral, antral, atretic, or lytic follicle) based on the transcriptional phenotype of mural granulosa cell within that phenotype as labeled during the multi-level cell type label assignment for spatial transcriptomics datasets.

### Single-cell regulatory network inference for Slide-seq datasets

pySCENIC implementation (v0.12.1) of the SCENIC pipeline^54,55^ (Single-Cell rEgulatory Network Inference and Clustering) was used to infer transcription factors, gene regulatory networks for Slide-seq data. Raw spatial gene expression counts for combined Slide-seq datasets were saved in loom file format using the loompy package (v2.0.17). These counts were used for gene regulatory network inference and generation of co-expression modules using pySCENIC package. Inferred modules were used for regulon prediction aka cisTarget and cellular enrichment of regulons (AUCell) was estimated for every bead in Slide-seq data. AUCell regolon enrichment scores were compared between representing different stages of luteinizing hormone treatment to estimate Regulon Specificity Scores (RSS) for specific stages/ conditions.

### Temporal gene expression analysis and clustering for Slide-seq datasets

Custom scripts were used to study temporal trends in gene expression across stages of preovulatory ovaries. In short, Generalized Additive Models (GAMs) were trained on expression values of highly variable genes across discrete time points using the pyGAM package^56^ (v0.8.0). LinearGAM function as implemented in pyGAM package was used with n_splines=4 and spine_order=2 to derive the GAM models for individual genes which were then used to predict gene expression trends on the same eight discrete time points. Predicted gene expression values for all genes were then scaled and clustered using K-means clustering with k from 4-6 as implemented in the scikit-learn^57^ package (v1.2.0). Scaled gene expression trends were then visualized for temporally variable genes within gene groups. Genes of interest within individual gene groups were visualized on spatial maps using scanpy. Pseudo-bulk granulosa mural cell transcriptomes at a follicle resolution were used to study temporal changes in mural granulosa cells. Raw gene expression counts for mural granulosa cells within each follicle were added to construct pseudo-bulk granulosa mural transcriptomes.

### Construction and analysis of pseudo-bulk oocyte transcriptomes

Bulk oocyte transcriptomes were constructed from spatial transcriptomics beads under individual oocyte cells. Beads classified as oocytes during deconvolution were extracted and filtered to remove beads with oocyte prediction score <= 0.5 and select the beads most likely to be in oocytes. Spatial distances between all pairs of beads under oocytes were then used to filter out beads with less than five beads within 100 µm distance. Individual samples were then reprocessed, and an additional round of filtering was performed to remove prediction noise. Raw gene expression counts for oocyte beads within each follicle were added to construct pseudo-bulk oocyte transcriptomes.

### Cell-cell interactions analysis on spatial transcriptomics data

Cell-cell signaling analysis was performed using the ligand-receptor interaction analysis module of squidpy package. For oocytes and cumulus cells in preovulatory follicles, mean expression of annotated ligand-receptor interaction pairs from Omnipath database was calculated at follicle resolution using the squidpy implementation of the method CellPhoneDB^58^. Mean expression values of ligand receptor paris across follicles were then compared between different stages of ovulation to identify stage specific cell-cell interactions. For interactions between dividing mesenchyme and granulosa cells in the immature untreated ovaries, ligand receptor interaction analysis on all annotated pairs was performed on beads classified as granulosa and dividing mesenchyme.

### Enrichment analysis for Gene Ontology (GO) for spatial transcriptomics

GO term enrichment analysis was performed on differentially expressed genes using gseapy^59,60^ (version v1.0.4) wrapper package. Differential gene expression analysis results were used to select genes which were used for GO term enrichment analysis using GO_Biological_Processes_2021 gene sets in enrichr command. The enriched GO terms of interest were selected and visualized on a bar plot. The differentially expressed genes that were associated with GO terms of interest were used to calculate module scores using the score_genes command in scanpy. Module scores for individual beads then compared across different follicle types and across stages of ovulation.

### Sample preparation for RNA fluorescence in-situ hybridization (FISH), immunofluorescence, and histology

Whole ovaries were isolated using aseptic technique and placed in ice cold sterile 1X PBS. Fresh tissues were immediately embedded in Optimal Cutting Compound (OCT) media and frozen in liquid nitrogen cooled isopentane, cut into 10 µm sections using a Thermo Scientific Microm 550 cryostat, and mounted on −20°C cooled histological glass slides which were then stored at −80°C until used.

### RNA fluorescence in-situ hybridization (FISH) split probe design and Signal Amplification using Hybridization Chain Reaction HCR-V3

Two-step hybridization strategy with split probe design and Hybridization Chain Reaction (HCR)-V3^61^ was used to label up to three transcripts in a single tissue section. Split probes for individual gene targets were designed using NCBI primer-blast^62^. The probes were checked for binding specificity against the RefSeq mRNA database for *Mus musculus*. We designed 20-21 bp primer pairs for an amplicon length of 40-42 bp (2 x primer length), primer melting temperature between 57°C and 63°C, and primer GC content between 35% and 65%. 5-10 pairs of reverse complemented forward primers and reverse primers were then concatenated to the flanking initiator sequence for HCR, ordered from Integrated DNA Technologies (IDT) with standard desalting purification. Split probes for each gene target, mixed and diluted in nuclease-free water to create a split probe pool stock solution at 10µM total probe concentration for every target. Hairpin pairs labeled with three different fluorophores namely Alexa-488, Alexa-546, and Alexa-647 (Molecular Instruments) were used for HCR-V3.

### RNA fluorescence in-situ hybridization (FISH) experiments

Slides with tissue sections were then brought to room temperature until the OCT melts and were then immediately fixed in 4% paraformaldehyde for 12 minutes at room temperature. Post fixation, the sections were washed for 5 mins in 1x PBS twice, incubated for 1 hour in 70% ethanol for tissue permeabilization, washed again for 5 mins in 1x PBS, and then used for primary hybridization. Hybridization Buffer (HB) mix was prepared with 2x SSC, 5x of Denhart Solution, 10% Ethylene Carbonate, 10% Dextran Sulfate, 0.01% SDS, 1µM of probe pool mix per target for the hybridization reaction. 20-30 µl of HB mix (with probes) per section was then put on each slide to cover the tissue section, covered with parafilm, and incubated overnight at 37°C inside a humidifying chamber for primary hybridization. After primary hybridization, parafilm was removed and slides were washed in Hybridization Wash Buffer-1 (0.215M NaCl, 0.02M Tris HCl pH 7.5, and 0.005M EDTA) for 20-30 minutes at 48°C. Amplification Buffer (AB) mix was prepared with 2x SSC, 5x of Denhart Solution, 10% Dextran Sulfate, 0.01% SDS, 0.06µM of HCR hairpins for the amplification reaction. 2µl of each fluorophore labeled hairpins at 3µM corresponding to the target genes were mixed, incubated at 95°C for 1.5 minutes, covered in aluminum foil, and left to cool down at room temperature for 30 minutes to form hairpins before adding it to AB mix. 20 µl of AB mix per section was then put on each slide to cover the tissue section, covered with parafilm, and incubated overnight at room temperature in the dark for signal amplification. After signal amplification, parafilm was removed, and slides were washed in the 5x SSCT buffer twice for 30-40 minutes and then twice for 10 mins. The slides were then carefully cleaned with Kimwipe and treated with Ready Probes Auto-fluorescence Quenching Reagent Mix (Thermo Fisher, R37630) for 5 minutes and washed for 5 minutes in 1X PBS twice. Last, tissue sections were counterstained with DAPI for 5 minutes at room temperature, washed for 5 minutes in 1x PBS twice, excess PBS cleaned using Kimwipe, immediately mounted on coverslips using Slowfade antifade media, left overnight for treatment, and imaged the next day on a Zeiss Axio Observer Z1 Microscope using a Hamamatsu ORCA Fusion Gen III Scientific CMOS camera. smFISH images were shading corrected, stitched, rotated, thresholded, and exported as TIFF files using Zen 3.1 software (Blue edition).

## DATA AVAILABILITY

The authors declare that all sequencing data supporting the findings of this study have been deposited in NCBI’s Gene Expression Omnibus (GEO) with GEO series accession number <u>GSE240271</u>. Bead barcode location files matched to spatial transcriptomics datasets have been made publicly available on GitHub (https://github.com/madhavmantri/mouse_ovulation). All other data supporting the findings in this study are included in the main article and associated files. Source data for individual figures are provided with this paper.

## CODE AVAILABILITY

Scripts to reproduce the analysis presented in this study have been deposited on GitHub (https://github.com/madhavmantri/mouse_ovulation).

### ACKNOWLEDGEMENTS

We would like to thank Dr. Peter Schweitzer and the Cornell Genomics Center for help with sequencing assays and the Cornell Bioinformatics facility for assistance with bioinformatics. We also thank the members of the Ren and De Vlaminck labs for many valuable discussions. This work was supported by a seed grant provided by Cornell Center of Vertebrate Genomics (to Y.A.R. and I.D.V.) and R01HD109392 from NICHD (to Y.A.R).

## AUTHOR CONTRIBUTIONS

M.M., Y.A.R., and I.D.V. designed the study. M.M. and H.H.Z. performed the animal experiments. M.M. and E.S. performed the spatial transcriptomics experiments. M.M. and E.S. analyzed the data. M.M., H.H.Z., and E.S. performed histology, RNA FISH, and analyzed the images. M.M., H.H.Z., E.S, Y.A.R., and I.D.V. wrote the manuscript. All authors provided feedback and comments.

## COMPETING INTERESTS

The authors declare no competing interests.

## Notes

https://www.ncbi.nlm.nih.gov/geo/query/acc.cgi?acc=GSE240271

### Competing Interest Statement

The authors have declared no competing interest.

### Summary of Updates

All images in main article and supplement files were replaced with higher resolution images. Line numbers were added to the main article PDF. Correction was made in the Initials of one of the author names in Author Contribution section of the manuscript.

