## Supplemental Information for "A Spatiotemporal Molecular Atlas of the Ovulating Mouse Ovary"

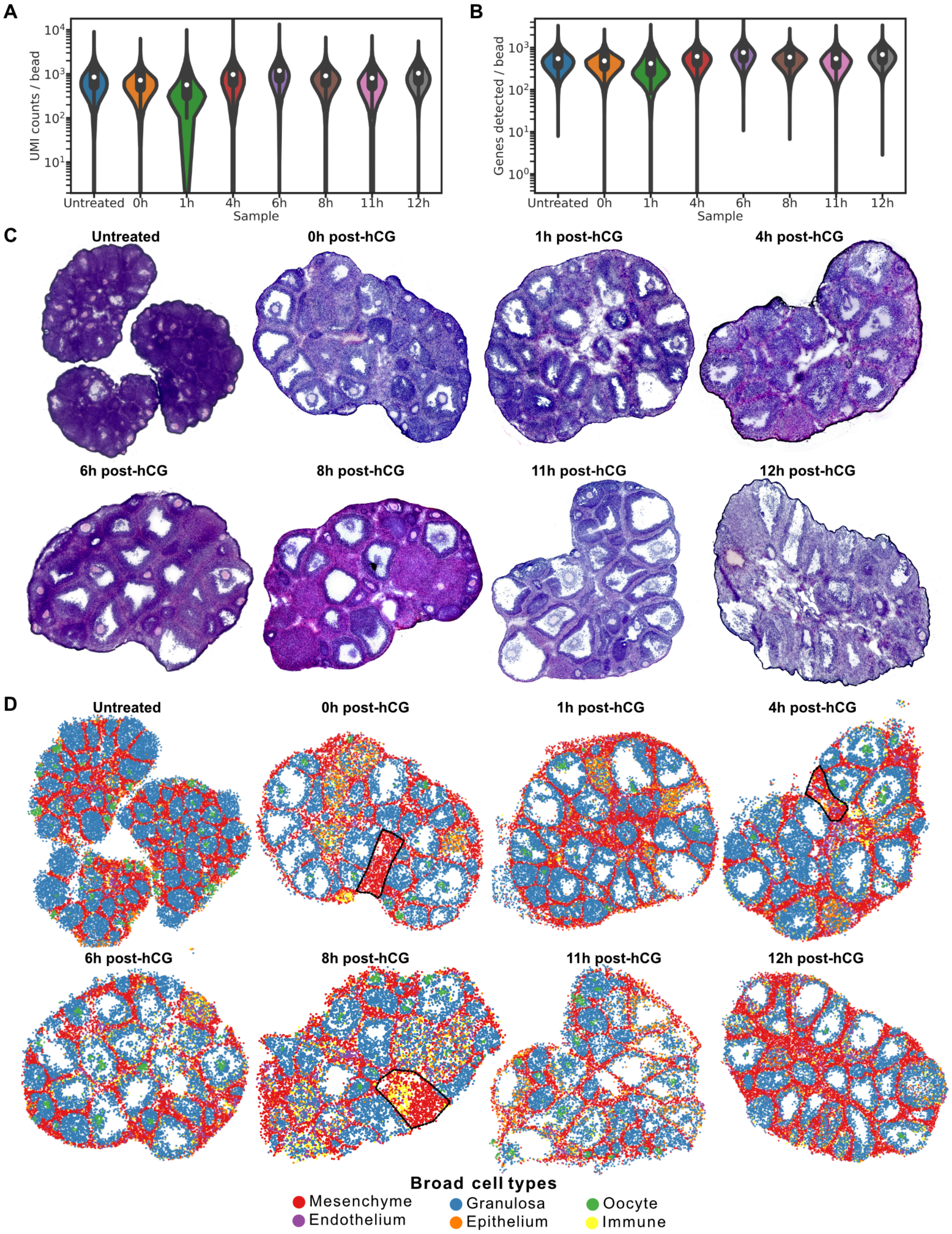


**Supplementary Figure 1**: **Spatial transcriptomics of hormone-induced ovulation. A)** Violin plots showing the number of unique molecular identifiers (UMIs) detected per bead across samples. **B)** Violin plots showing the number of genes detected per bead across samples. **C)** Images of Hematoxylin and Eosin-stained ovary tissue sections consecutive to the sections used for spatial transcriptomics experiments. **D)** Spatial transcriptomics maps of three immature and seven preovulatory ovaries colored by broad cell type labels as predicted by deconvolution performed using scRNA-seq as a reference.

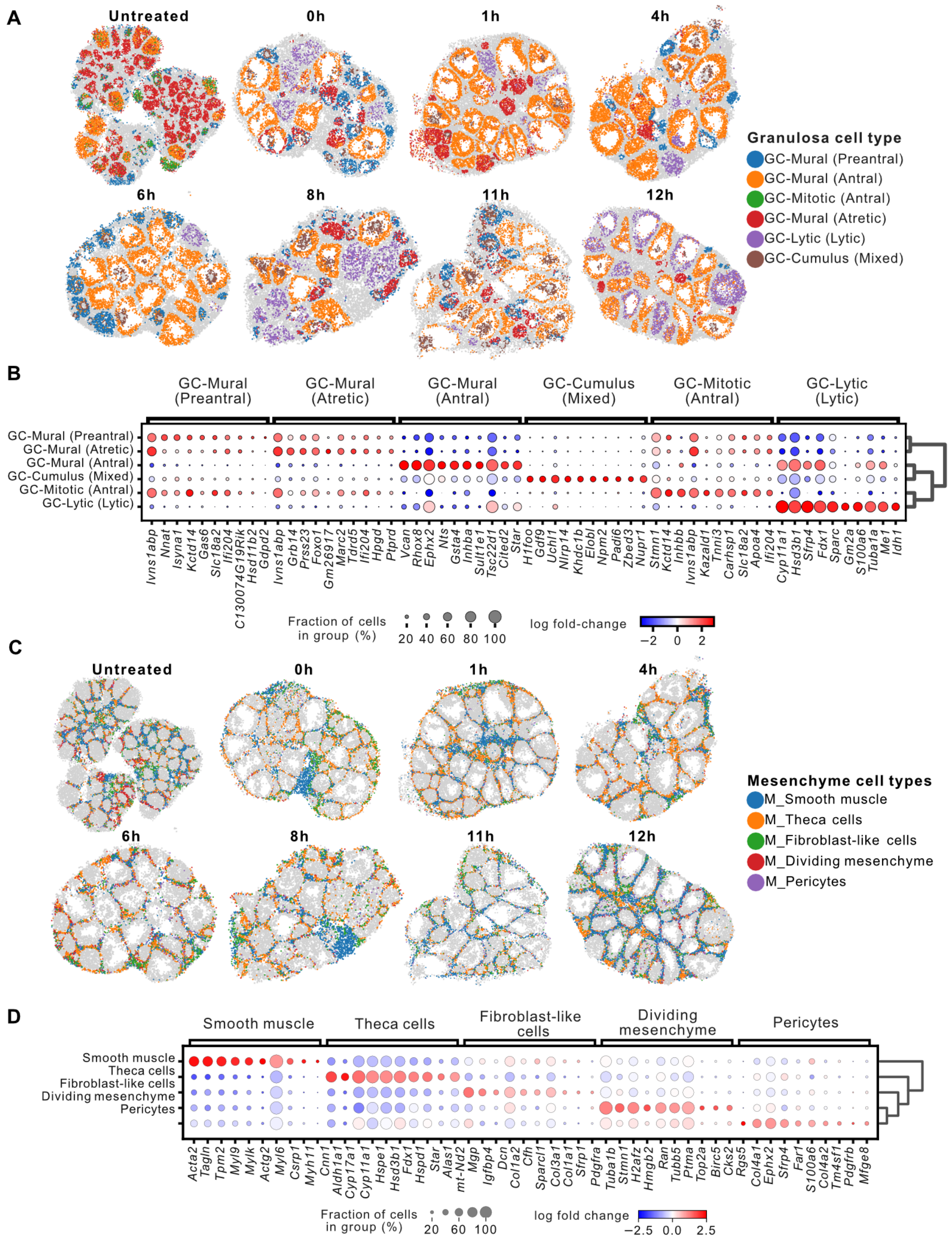


**Supplementary Figure 2**: **Cell type classification. A)** Spatial transcriptomics maps of three immature and seven preovulatory ovaries colored by fine cell type labels of granulosa cells, assigned using spatial location in conjugation with canonical markers of follicle states. **B)** Dot plot showing differential gene expression results for distinct granulosa cell types captured across three immature and seven preovulatory ovaries. **C)** Spatial transcriptomics maps of three immature and seven preovulatory ovaries colored by fine cell type labels of mesenchymal cells, as predicted by deconvolution performed using scRNA-seq as a reference. **D)** Dot plot showing differential gene expression results for distinct mesenchymal cell types captured across three immature and seven preovulatory ovaries.

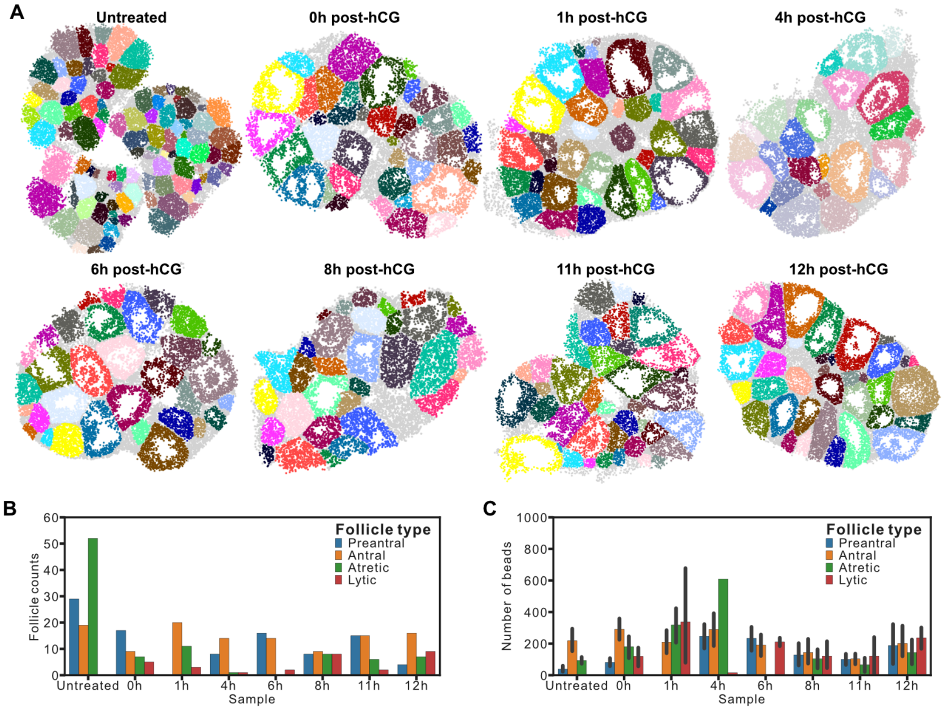


**Supplementary Figure 3**: **Temporal trends of follicle state. A)** Spatial transcriptomics maps of three immature and seven preovulatory ovaries with 350 follicles manually labeled. **B)** Bar plot showing the number of each follicle type present across eight time points before and after hormone treatment. **C)** Bar plot showing the number of beads corresponding to follicle type across eight time points before and after hormone treatment.

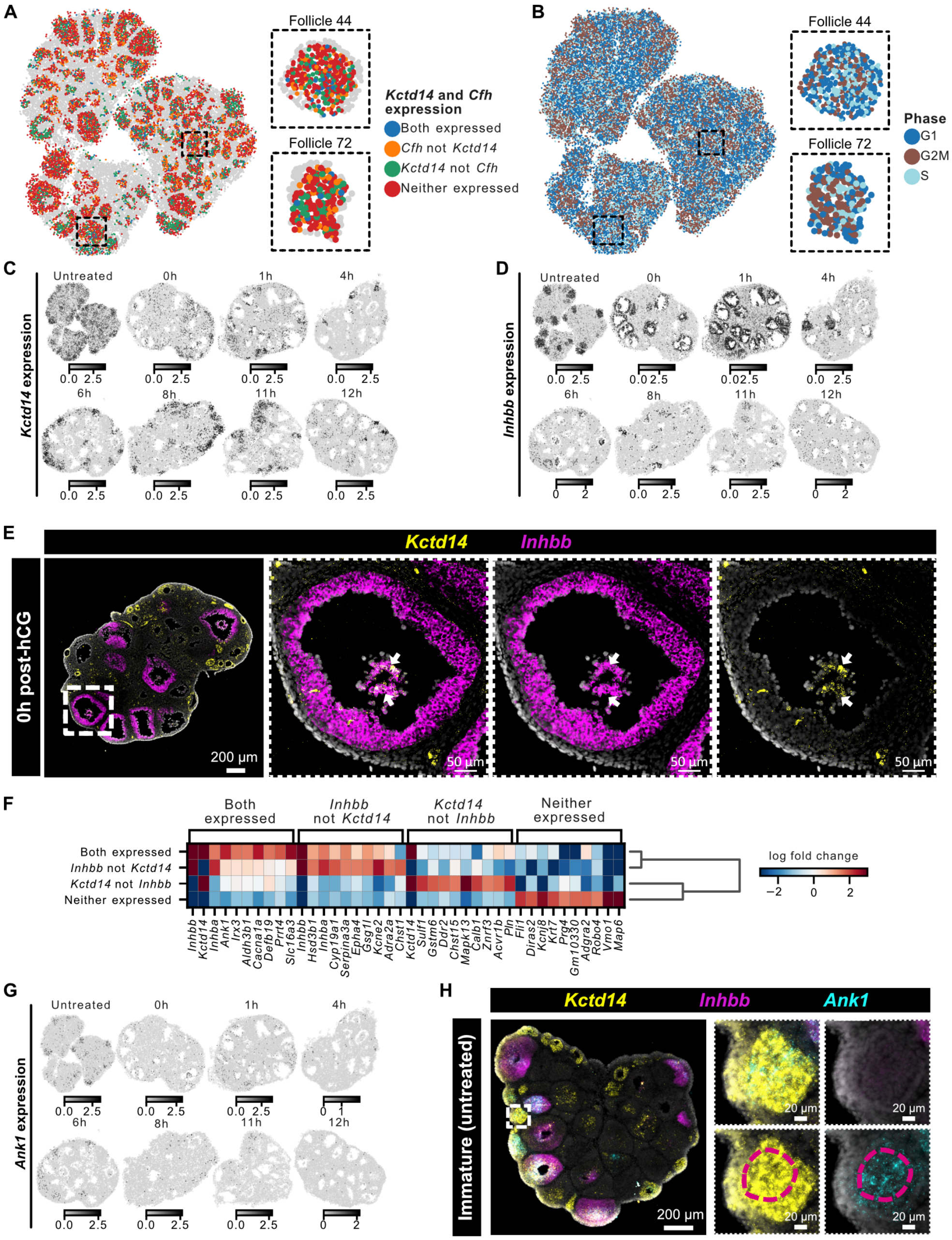


**Supplementary Figure 4**: **Spatial gene expression of granulosa cell markers associated with follicle state in immature ovaries. A)** Spatial transcriptomics maps of three immature ovaries showing *Kctd14* and *Cfh* expression. Two representative atretic follicles are highlighted in insets. **B)** Spatial transcriptomics maps of three immature ovaries showing cell cycle phase. Two representative atretic follicles are highlighted in insets. **C)** Spatial transcriptomics maps showing expression of *Kctd14*, a gene specific to preantral follicle phenotype. **D)** Spatial transcriptomics maps showing expression of *Inhbb*, a gene specific to antral follicle phenotype. **E)** Multiplexed RNA FISH staining for preantral marker *Kctd14* (yellow), and antral marker *Inhbb* (magenta) on tissue section from whole ovary collected at 0h post-hCG. The dotted boxes show zoomed-in images of a representative preovulatory follicle. Arrows point to *Kctd14*+*Inhbb*+ cumulus cells in preovulatory follicles. **F)** Heatmap showing differentially expressed genes in mural granulosa cells grouped by *Kctd14* and *Inhbb* expression. **G)** Spatial transcriptomics maps showing expression of *Ank1* (left) in three immature untreated ovaries. **H)** Multiplexed RNA FISH staining for preantral marker *Kctd14* (yellow), antral marker *Inhbb* (magenta), and *Ank1* gene (cyan) on untreated immature ovary. The dotted red regions show *Ank1*+ cells in a representative preantral follicle.

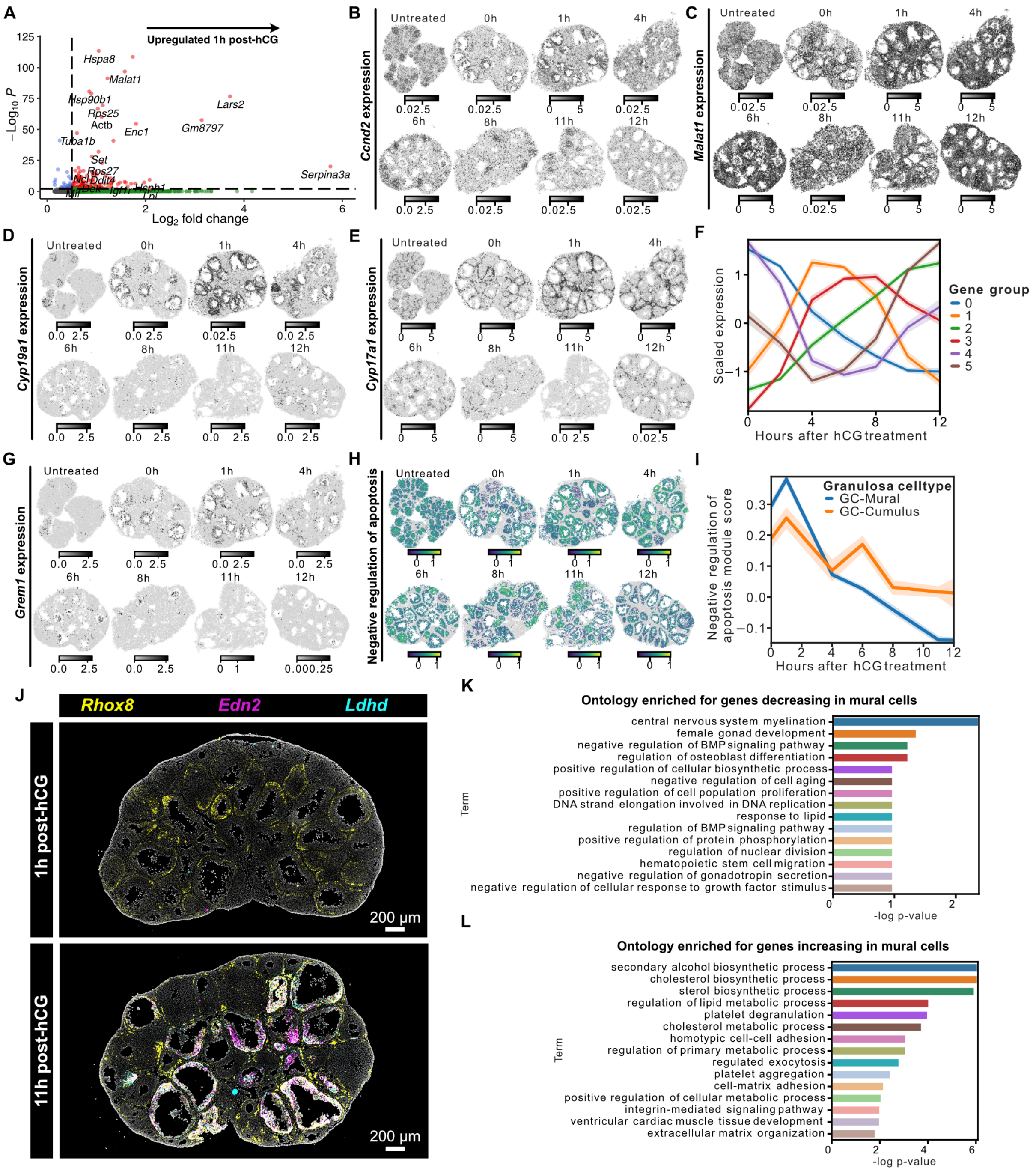


**Supplementary Figure 5**: **Analysis of mural granulosa cells post hormone treatment. A)** Volcano plot showing differentially expressed genes between 0h and 1h post hormone surge computed using two-sided Wilcoxon Rank-Sum test. Dotted lines represent thresholds for significantly enriched genes (red): -$\log_2$ fold change > 0.5 and p-value < $10^{-3}$. **B)** Spatial transcriptomics maps showing expression of *Ccnd2* gene in immature and preovulatory ovaries. **C)** Spatial transcriptomics maps showing expression of *Malat1* gene in immature and preovulatory ovaries. **D)** Spatial transcriptomics maps showing expression of *Cyp19a1* gene in immature and preovulatory ovaries. **E)** Spatial transcriptomics maps showing expression of *Cyp17a1* gene in immature and preovulatory ovaries. **F)** Line plot showing gene expression trends for groups of genes clustered by their temporal expression in mural granulosa cells post hormone treatment. **G)** Spatial transcriptomics maps showing expression of *Grem1* gene in immature and preovulatory ovaries. **H)** Spatial transcriptomics maps showing gene module score for negative regulation of apoptosis in all granulosa cells. **I)** Line plot showing a comparison of dynamics of gene module score for negative regulation of apoptosis between mural and cumulus granulosa cells in preovulatory follicles across eight time points in immature and preovulatory ovaries. **J)** Multiplexed RNA FISH staining for granulosa cell marker *Rhox8* (yellow), rupture-associated marker *Edn2* (magenta), and metabolic activity gene *Ldhd* (cyan) on tissue sections from whole ovaries collected at 1h and 11h post-hCG. Representative images from four biological replicates. **K)** Bar plot showing top 15 gene ontology (GO) terms for genes decreasing in mural granulosa cells after hCG treatment. **L)** Bar plot showing top 15 gene ontology (GO) terms for genes increasing in mural granulosa cells after hCG treatment.

---

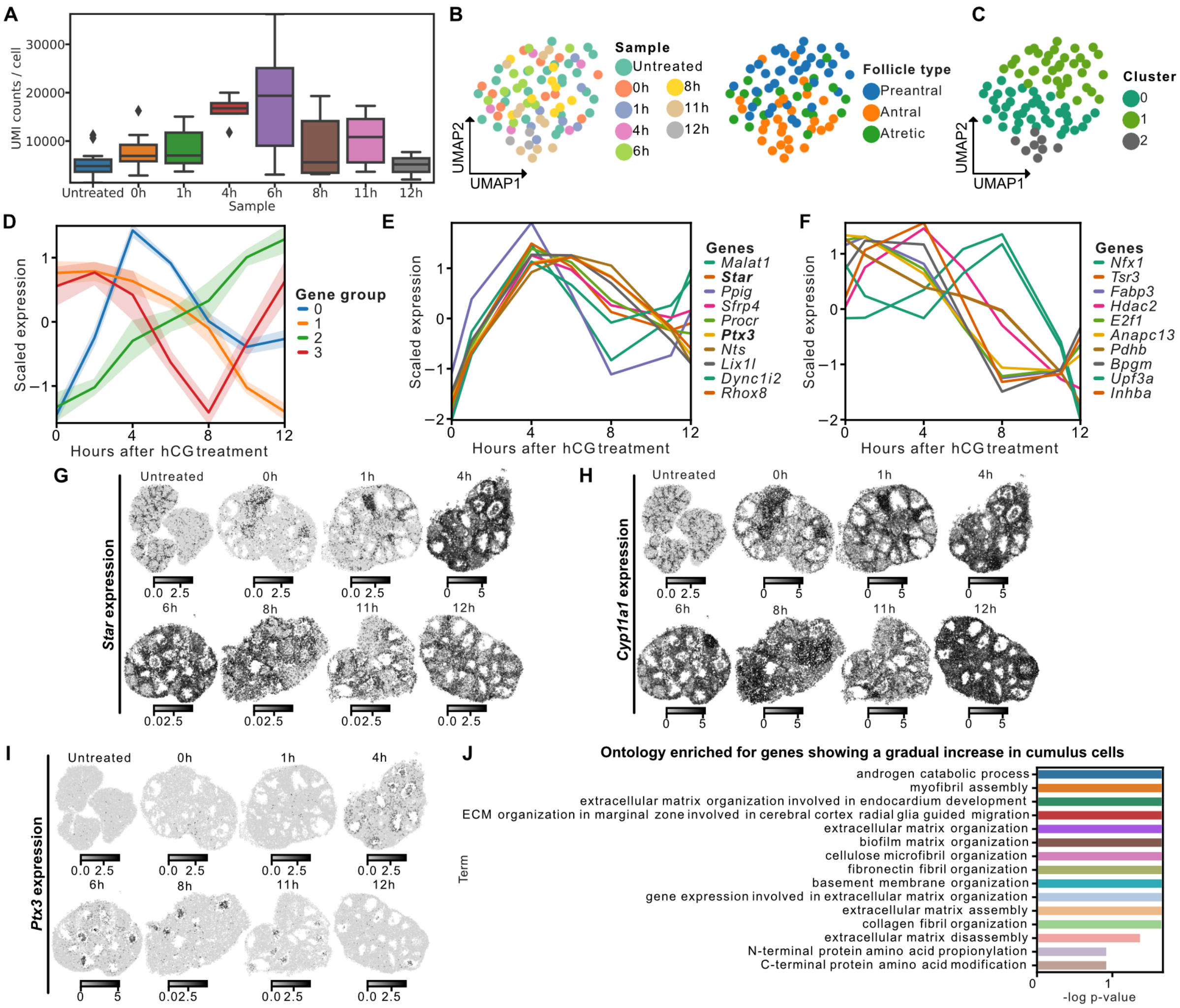


**Supplementary Figure 6**: **Analysis of oocyte and cumulus cells post hormone treatment. A)** Box plot showing the number of unique molecular identifiers (UMIs) detected per oocyte cell across samples. **B)** UMAP showing oocyte transcriptomes clustered by pseudo-bulk gene expression and colored by sample (left), and follicle type (right). **C)** UMAP showing oocyte transcriptomes clustered by pseudo-bulk gene expression and colored by oocyte cluster ID. **D)** Line plot showing gene expression trends for groups of genes clustered by their temporal expression in cumulus cells post hormone treatment. **E)** Line plot showing temporal trends of genes that show a sharp increase in expression between 0h and 4h post-hormone treatment in cumulus cells from preovulatory follicles. **F)** Line plot showing temporal trends of genes that show a decrease in expression with time post hormone surge in cumulus cells from preovulatory follicles. **G)** Spatial transcriptomics maps showing expression of *Star*, a gene that encodes a protein involved in regulation of steroid hormone synthesis. **H)** Spatial transcriptomics maps showing expression of *Cyp11a1*, a gene that encodes a protein involved in the synthesis and metabolism of cholesterol, steroids and other lipids. **I)** Spatial transcriptomics maps showing expression of *Ptx3*, a gene that encodes a major component of the extracellular matrix of the cumulus-oocyte complex during cumulus expansion. **J)** Bar plot showing top 15 gene ontology (GO) terms for genes showing a gradual increase in cumulus cells after hCG treatment.

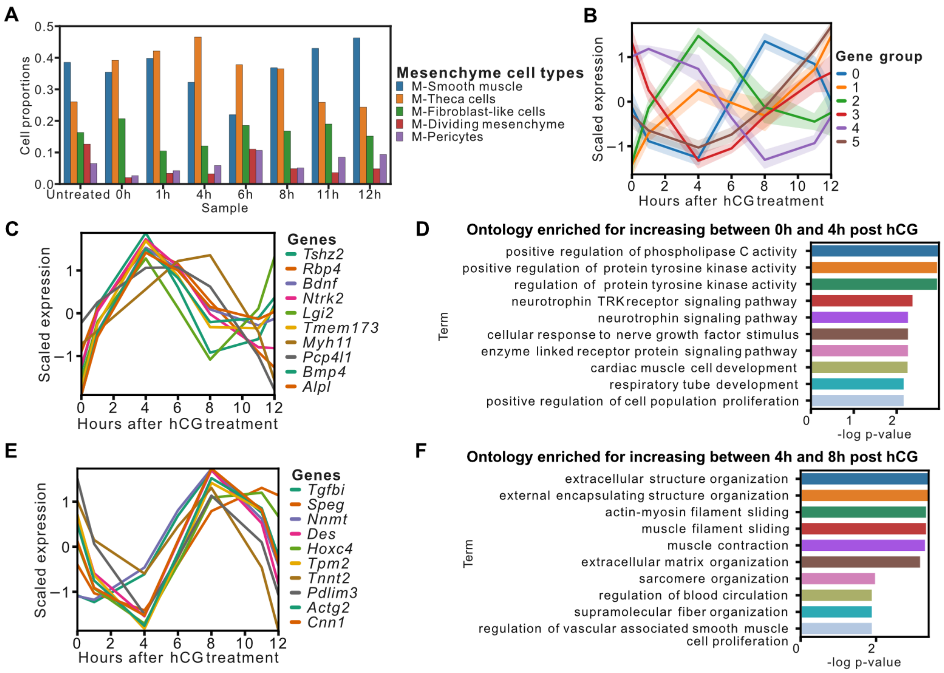


**Supplementary Figure 7**: **Analysis of stromal cells post hormone treatment. A)** Bar plot showing proportions of mesenchyme-specific cell types across all time points. **B)** Line plot showing gene expression trends for groups of genes clustered by their temporal expression in mesenchymal cells post hormone treatment. **C)** Line plot showing temporal trends of genes that show a sharp increase in expression between 0h and 4h post-hormone treatment in mesenchyme cells from preovulatory follicles. **D)** Bar plot showing top 10 gene ontology (GO) terms for genes showing a sharp increase in mesenchymal cells between 0h and 4h after hCG treatment. **E)** Line plot showing temporal trends of genes that show a sharp increase in expression between 4h and 8h post-hormone treatment in mesenchyme cells from preovulatory follicles. **F)** Bar plot showing top 10 gene ontology (GO) terms for genes showing a sharp increase in mesenchymal cells between 4h and 8h after hCG treatment.

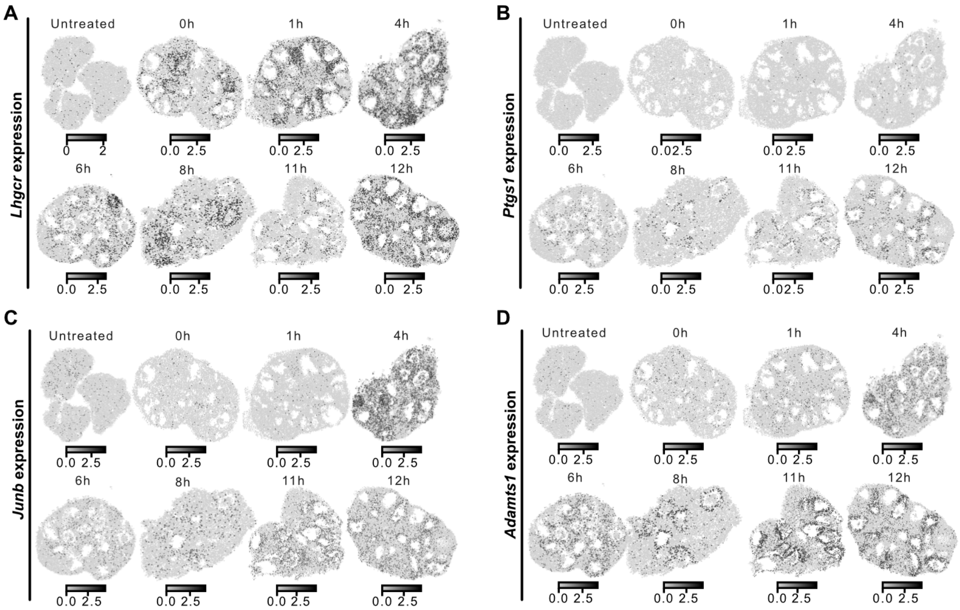


**Supplementary Figure 8**: **Follicle heterogeneity before ovulation. A)** Spatial transcriptomics maps showing expression of *Lhgcr* gene in immature and preovulatory ovaries. **B)** Spatial transcriptomics maps showing expression of *Ptgs1* gene in immature and preovulatory ovaries. **C)** Spatial transcriptomics maps showing expression of *Junb* gene in immature and preovulatory ovaries. **D)** Spatial transcriptomics maps showing expression of *Adamts1* gene in immature and preovulatory ovaries.
